# Differential Effects of 5-HT7 Receptor Signaling in the Noradrenergic System in a Rat Model of Treatment-Resistant Depression

**DOI:** 10.64898/2026.01.05.697635

**Authors:** C Bruzos-Cidón, N Llamosas, H Bengoetxea, JV Lafuente, L Ugedo, M Torrecilla

## Abstract

**Introduction:** Treatment-resistant depression (TRD) is a major clinical challenge, and its neurobiological basis remains unclear. The locus coeruleus (LC), a key noradrenergic hub, shows altered activity in TRD models, and serotonergic 5-HT_7_ receptors have emerged as potential modulators of this circuit.

**Methods:** We assessed the impact of 5-HT_7_ receptor activation on LC function in Wistar Kyoto rats, an established TRD model, using in vivo extracellular recordings, in vitro patch-clamp assays, and receptor expression analysis.

**Results:** Systemic administration of AS19, a selective 5-HT_7_ receptor agonist, increased LC neuronal firing in both strains, with a significantly greater effect in Wistar Kyoto rats. This response was prevented by a 5-HT_7_ antagonist confirming receptor specificity. Glutamatergic blockade altered AS19 effects in a strain-dependent manner, whereas in vitro assays revealed no differential postsynaptic modulation and similar presynaptic inhibition across groups. Western blot analysis showed elevated 5-HT_7_ receptor expression in the LC, ventral hippocampus, and amygdala of Wistar Kyoto rats, alongside reduced expression in the prefrontal cortex.

**Conclusion:** These findings indicate that 5-HT_7_ receptor activation exerts complex, circuit-dependent modulation of LC activity, influenced by glutamatergic and serotonergic inputs. Enhanced receptor expression in limbic regions and altered LC responsiveness may contribute to TRD pathophysiology and highlight 5-HT_7_ signaling as a promising target for novel therapeutic strategies.

## 1 Introduction

Major Depressive Disorder (MDD) is highly prevalent and a major global health burden with substantial economic costs. According to the World Health Organization, MDD is the leading cause of healthy life years lost worldwide, worsened by the COVID-19 pandemic (World Health Organization, 2022). Despite multiple therapeutic options, many patients report residual symptoms (McClintock et al., 2011; Zajecka et al., 2013), and some fail to respond to antidepressants, a condition called treatment-resistant depression (TRD) (McIntyre et al., 2023). Animal models have helped clarify depression neurobiology and mechanisms of current and potential treatments (Millan et al., 2003; Willner, 2017).

The Wistar Kyoto (WKY) rat is a well-established depression model (Aleksandrova et al., 2019; Millard et al., 2020). This strain shows depressive-like behaviors (Lahmame & Armario, 1996; Lahmame et al., 1997; López-Rubalcava & Lucki, 2000; Nam et al., 2014; Tejani-Butt et al., 2003; Will et al., 2003) and resistance to SSRIs and other monoaminergic drugs (Holl et al., 2018; Kin et al., 2019; López-Rubalcava & Lucki, 2000; Smith et al., 2019), making it suitable for TRD research. Dysregulation of the noradrenergic system is linked to the WKY phenotype (Conti et al., 1997; Ma & Morilak, 2004; Pardon et al., 2002; Pardon et al., 2003; Paré & Tejani-Butt, 1996; Sands et al., 2000; Tejani-Butt et al., 1994; Zafar et al., 1997). Functional abnormalities occur in the locus coeruleus (LC), the main source of central noradrenaline (NE). WKY rats exhibit reduced basal NE levels (De La Garza & Mahoney, 2004; Scholl et al., 2010; Yamada et al., 2013) and increased expression of genes for NE turnover (Pearson et al., 2006), and functionally, higher LC firing and burst activity with diminished inhibitory control via α_2_-adrenoceptors and GABAergic inputs (Bruzos-Cidón et al., 2015; Bruzos-Cidón et al., 2014).

Given its role in depression, the LC is a key target for studying antidepressant mechanisms, including vortioxetine (Pehrson et al., 2013), a multimodal drug approved for MDD (Gibb & Deeks, 2014). Vortioxetine inhibits serotonin transporters and modulates several receptors, notably 5-HT_7_ (Bang-Andersen et al., 2011), which has the highest serotonin affinity (Ruat et al., 1993). This Gs/G_12_-coupled receptor is widely expressed in rodent brain regions such as hypothalamus (HPT), thalamus, hippocampus (HPC), cortex, amygdala (A), striatum, spinal cord, cerebellum, and dorsal raphe nucleus (DRN) (Bonaventure et al., 2004; Hedlund & Sutcliffe, 2004; Neumaier et al., 2001). Autoradiographic studies indicate a similar distribution in the human brain, with high expression in thalamus, DRN, HPC, and HPT (Martín-Cora & Pazos, 2004; Varnäs et al., 2004).

Preclinical evidence supports 5-HT_7_ receptors as promising antidepressant targets. Antagonism produces antidepressant-like effects in rodents (Abbas et al., 2009; Hedlund et al., 2005; Mnie-Filali et al., 2011; Wesołowska et al., 2006), and knockout mice lacking 5-HT_7_ receptors show similar phenotypes (Cates et al., 2013; Guscott et al., 2005). Chronic SB-269970 reverses depressive-like behavior and desensitizes 5-HT_1A_ autoreceptors (Mnie-Filali et al., 2011). SB-269970 also mitigates synaptic plasticity deficits induced by maternal fluoxetine exposure (Bobula et al., 2024).

Conversely, 5-HT_7_ overexpression in the developing prefrontal cortex (PFC) increases glutamatergic input to the DRN and promotes depressive-like behaviors (Olusakin et al., 2020). Activation of 5-HT_7_ receptors influences hippocampal dendritic spine remodeling and depressive-like behavior (Bijata et al., 2022) and modulates DRN activity via GABAergic mechanisms (Kusek et al., 2015; Roberts et al., 2001; Roberts et al., 2004). AS19, a selective 5-HT_7_ agonist widely used in preclinical studies (Leopoldo, 2004), helps investigate functional impact in relevant circuits. Reciprocal interactions between DRN and LC regulate neuronal firing and neurotransmitter dynamics (Pudovkina et al., 2002), making LC a key target for understanding serotonergic modulation.

Given evidence linking 5-HT_7_ signaling to depression and antidepressant mechanisms, this study aimed to assess the impact of 5-HT_7_ activation on LC neurophysiology in WKY rats. Specifically, we examined whether 5-HT_7_-mediated effects differ between WKY and the control strain, the Wistar rat (Wis) using in vivo extracellular and in vitro patch-clamp recordings in LC slices.

## 2 Material and Methods

### 2.1 Animals

Male Wis (n = 32) and WKY (n = 29) rats weighting 175–300 g were used for the in vivo and in vitro electrophysiological recordings, and western blot experiments. This weight range ensures the use of age-matched animals, according to the animal supplier. Every effort was made to minimize the suffering of the animals and to use the minimum number of animals possible. Experimental protocols were reviewed and approved by the Local Committee for Animal Experimentation at the University of the Basque Country. All the experiments were performed in compliance with the European Community Council Directive on ‘The Protection of Animals Used for Experimental and Other Scientific Purposes’ (2010/63/UE) and with the Spanish Law (RD 53/2013) for the care and use of laboratory animals.

### 2.2 Drugs

(2S)-N,N-dimethyl-5-(1,3,5-trimethylpyrazol-4-yl)-1,2,3,4-tetrahydronaphthalen-2-amine (AS19), 4-Hydroxyquinoline-2-carboxylic acid (kynurenic acid), (2R)-1-[(3-Hydroxyphenyl) sulfonyl]-2-[2-(4-methyl-1-piperidinyl) ethyl] pyrrolidine hydrochloride (SB269970 hydrochloride) and picrotoxin were purchased from Tocris Bioscience, UK. Chloral hydrate from Sigma-Aldrich, USA and (S)-1-Aminopropane-1,3-dicarboxylic- acid (L-glutamic acid) from Research Biochemicals International, USA.

AS19 for in vivo administration was prepared in 5% polyethylene glycol solution and SB 269970 hydrochloride was prepared in distilled water. For in vitro recordings, all drugs were prepared in distilled water every day and then diluted in artificial cerebrospinal fluid (aCSF). Chloral hydrate was prepared in 0.9% saline andKynurenic acid for intracerebroventricular (i.c.v.) injections in Dulbecco (in mM): NaCl 136.9; KCl 2.7; NaH2PO4 8.1; KH2PO4 1.5; MgCl2 0.5; CaCl2 0.9. It was firstly dissolved in NaoH 1N and pH-adjusted to 7.4. Finally, an appropriate volume of sterile 45 mM CaCl₂ stock was added at the end to achieve the desired concentration while minimizing the risk of calcium precipitation.

### 2.3 Electrophysiological procedures

#### 2.3.1. Single-unit extracellular recording of locus coeruleus neurons in vivo

Single-unit extracellular recordings of LC were performed as previously described (Bruzos-Cidón et al., 2015). Animals were anesthetized with chloral hydrate (400 mg/kg i.p.) and placed in a stereotaxic frame.

For LC recordings, the head was oriented at 15° to the horizontal plane (nose down). The recording electrode was lowered into the LC (relative to lambda: AP -3.7 mm, ML +1.1 mm, and DV -5.5 to - 6.5 mm). LC neurons were identified by standard criteria which included: spontaneous activity displaying a regular rhythm and a firing rate between 0.5–5 Hz; characteristic spikes with a long-lasting, positive-negative waveform, and a biphasic excitation–inhibition response to pressure applied to the contralateral hindpaw (paw pinch), as previously described (Pineda et al., 1996).

Firing pattern was analyzed offline, using the computer software Spike2 (Cambridge Electronic Design, UK) and the following parameters were calculated: firing rate (number of spikes in 10s) and the response to intravenous drug administration. Changes in firing rate are expressed as percentages of the baseline firing rate, which was taken as the mean firing rate during 3 min prior to drug injection. Only one cell was studied in each animal when any drug was administered.

#### 2.3.2. Patch-clamp recordings of locus coeruleus neurons from rat brain slices

Slice preparation and whole-cell patch-clamp recordings were performed as described by Bruzos-Cidón et al.(2015).

##### 2.3.2.1. Brain slice preparation

Briefly, animals were anesthetized with chloral hydrate (400 mg/kg, i.p) and decapitated. The brain was immediately extracted and placed in cooled aCSF containing (in mM): 125 NaCl, 2.5 KCl, 1.2 MgCl2, 26 NaH2CO3, 1.25 NaH2PO4, 2.4 CaCl2, 11 D-glucose saturated with 95% O2 and 5% CO2 (pH 7.3-7.4). During the incubation period, (+)-MK-801 maleate (10 μM) was added to aCSF in order to reduce neuronal damage. Horizontal brainstem sections (220 μm) containing the LC were cut using a vibrotome (VT1200S; Leica Microsystems, Germany). LC slices were incubated in warmed (35°C) aCSF for at least 30 min before recording.

##### 2.3.2.2. Neuronal identification and recording

The slice mounted on a recording chamber was maintained at 35-37°C and perfused with aCSF at a flow rate of 1.5-2 ml/min by gravity. LC neurons were visualized using an upright microscope with infrared optics (Eclipse E600FN, Nikon®) as a dense and compact group of cells located on the lateral border of the central gray and the fourth ventricle, just anterior to the genu of the facial nucleus in rat horizontal slices.

LC neurons were identified by the presence of an inwardly rectifying potassium current detected by typical current-voltage (I/V) relationship observed when the membrane potential was hyperpolarized from -50mV to -130 mV in -10 mV increments (100 msec per step)(Williams et al., 1982). Membrane potential of LC neurons was held at -60 mV.

The patch-clamp procedure in voltage-clamp mode (whole-cell configuration) was employed to measure spontaneous synaptic activity and steady whole-cell currents. For that purpose, pipettes (2-4 MΏ) prepared from borosilicate glass capillaries (World Precison Instrumets, UK) with a micropipette puller (PC-10, Narishige CO., LTD, Japan) were filled with internal solution (in mM): 70 CsSO4, 20 CsCl, 20 NaCl, 1.5 MgCl2, 5 HEPES, 1 EGTA, 2 Mg ATP, and 0.5 Na-GTP (pH: 7.4, 280 mOsm). To specifically evaluate spontaneous Excitatory Postsynaptic Currents (sEPSC) voltage-clamp recordings were performed with 100 µM picrotoxin in the aCSF to block GABAA receptors. Recordings were detected with an Axopatch-200B (Axon Instruments, Foster City, CA), filtered at 5 kHz and digitized with a Digidata 1322A (Axon Instruments, Foster City, CA), and stored on a personal computer for offline analysis.

Data acquisitions and analyses were performed using pClamp 10.3 software package (Clampex and Clampfit programs; Molecular Devices). Recordings were post hoc filtered at 1 kHz. Template detection was used to select spontaneous synaptic events with deflections < 7 pA excluded from analysis. The template was generated from the average of multiple spontaneous events, and the selection was fitted to 3.5 threshold of the template. Spontaneous synaptic activity from each neuron was measured from a one-minute period, and several parameters were analyzed before and after AS19 application: Interevent interval, amplitude, decay time, rise time and half-width of sEPSC (that is, event’s width at 50% of the peak amplitude). After running the detection protocol each peak was visually inspected.

### 2.4. Histological procedure

#### 2.4.1. Western blot

Relative expression levels of the 5-HTR_7_ receptor (rabbit anti-5HTR_7_; 1:500; Ref: PA1-41122; Thermo Fisher Scientific) was quantified in Hypothalamus (HPT), dorsal Hippocampus (HPCd), ventral Hippocampus (HPCv), prefrontal cortex (PFC), Amygadala, Periacueductal Gray (PGA) and LC.

Tissue was homogenized in RIPA lysis buffer, and equal amounts of proteins were separated on SDS-polyacrylamide gels (Bio-Rad, Hercules, CA, USA). Proteins were transferred onto PVDF membranes by semi-dry transfer in Trans-Blot® Turbo™ Transfer System (Bio-Rad). Membranes were incubated for 1 h in TBS buffer (100 mM Tris-HCl; 0.9% NaCl, 1% Tween 20, pH 7.4) containing 5% non-fat dry milk to block nonspecific binding sites. Blots were then incubated in primary antibodies overnight. Actin was used as a loading control (mouse anti-actin; 1:2000; Ref: A-2066, Sigma-Aldrich, St. Louis, USA). The following day, HRP conjugated anti-rabbit secondary antibody (1:20.000; Ref: A-6164; Sigma-Aldrich, Spain) or anti-mouse secondary antibody (1:20.000; Ref: A-9044; Sigma-Aldrich, Spain) was added for 1 h at room temperature, and a chemiluminescent detection system (Ref: 34076, SuperSignal® West Dura Extended Duration Substrate, Fisher Scientific, Spain) was used to visualize the immunoreactive proteins. Images were acquired using ChemiDoc™ XRS+ Imaging System (Bio-Rad, Hercules, CA, USA) and optical densities were quantified with Image Studio Digits 3.1 (LI-COR BioScience Biotechnology, Germany). Values are expressed as the relative change from the Wistar rats.

### 2.5. Statistical analysis of data

Data analyses were conducted in GraphPad Prism (v.9.01; GraphPad Software, Inc). Saphiro -Wilk normality tests were applied to determine data distributions, and the appropriate parametric or nonparametric statistical test was performed accordingly. Two-sided Student’s t-test was used for comparisons of spontaneous or drug induced changes in firing rate and drug effects on inward currents and sEPSC. Two-sample Kolmogorov Smirnov test (KS) was applied to compare frequency distribution of sEPSC frequency and amplitude before and during AS19. One-sample Student’s t-test was used to compare 5-HT_7_ expression between strains. Repeated two-way ANOVA followed by Šídák’s post-hoc was used to analyze the in vivo AS19 effect in WKY compared to Wis. Data throughout the text are presented as mean ± S.E.M and median. Differences were considered to be significant for p < 0.05.

## 3. Results

### 3.1. WKY LC neurons show greater response to 5-HT_7_ receptor stimulation in vivo

Here we evaluated the effect of 5-HT_7_ receptor stimulation in the firing rate of LC neurons from an animal model of depression, the WKY rat. For that purpose, we systemically injected increasing doses of the 5-HT_7_ receptor agonist, AS19 (1.25-10 mg/kg i.v.) and recorded the spontaneous firing rate of LC neurons from WKY (n = 6) and Wis rats (n = 10) (Fig. 1a-b). Repeated two-way ANOVA revealed that cumulative doses of AS19 increased the firing rate of LC neurons in a dose-dependent manner in both strains (Dose: F_(3, 39)_ = 52.73; p < 0.0001; Strain: F_(1, 14)_ = 5.22; p < 0.05). However, the effect of AS19 at the highest administered dose was greater in LC neurons of WKY compared to that in Wis rats (p < 0.01, Šídák’s multiple comparisons test; Fig. 1b).

**Figure 1.**
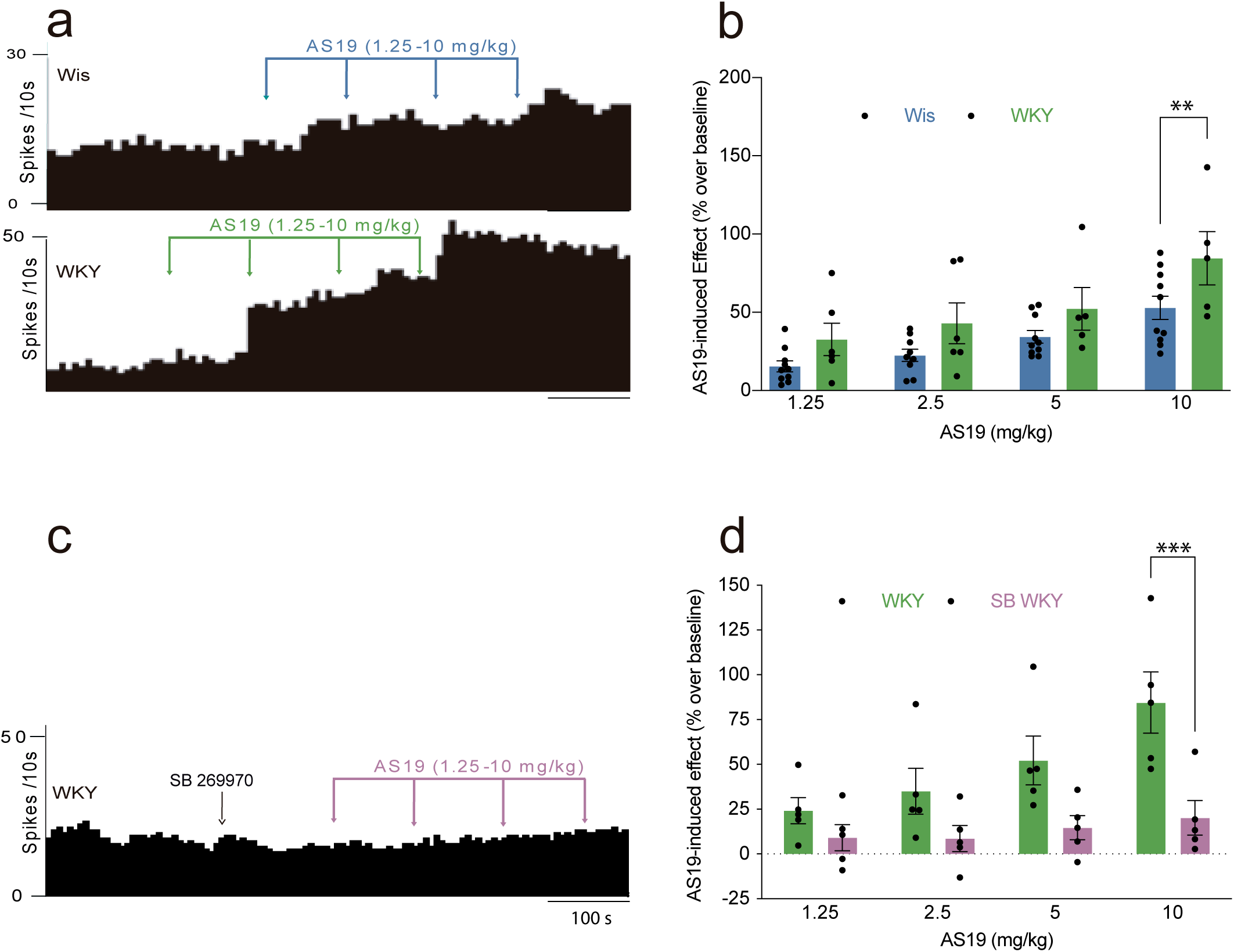
The stimulatory effect of AS19 on the basal firing rate of locus coeruleus (LC) noradrenergic neurons in Wistar (Wis) and Wistar Kyoto (WKY) rats. (a) Representative firing rate histograms illustrate the stimulatory effect of AS19 (1.25–10 mg/kg, i.v.) on LC basal activity in Wis and WKY rats. (b) Bar graphs showing AS19-induced effect in Wis (n = 10) and WKY (n = 6) rats. (**p < 0.01, Šídák’s multiple comparisons test). (c) Representative firing rate histogram represents the reduction of the AS19 effect produced by the administration of SB 269970 (1mg/kg, i.v.), the 5-HT_7_ receptor antagonist, on LC neurons in WKY (n = 4) rats. (d) Bar graphs showing statistically significant differences with the administration of 10 mg of AS19 on SB 269970 (1 mg/kg, i.v.) blockade in WKY r(n=4) rats. (***p < 0.001, Šídák’s multiple comparisons test). Bars represent mean ± S.E.M and circles individual cell values.

Next, we checked if the excitatory effect induced by AS19 was due to 5-HT_7_ receptor activation. For that purpose, SB 269970 (1 mg/kg i.v.), which has been described as a 5-HT_7_ receptor antagonist (Lovell et al., 2000) was injected before AS19 administration in WKY rats. Repeated measures two-way ANOVA revealed that SB 269970 reduced AS19 effect on LC neuron activity from WKY rats (AS19 x SB: F_(3, 24)_ = 13.31; p < 0.0001), which was statistically significant for the highest dose of AS19 (p < 0.001, Šídák’s multiple comparisons test) (Fig. 1c-d).

### 3.2. Differential glutamatergic modulation of the in vivo AS19 stimulatory effect in LC neurons of WKY rats

To study the potential contribution of glutamatergic transmission to the stimulatory effect of AS19 on LC neurons we administered intracerebroventricularly the non-selective glutamate receptor antagonist, kynurenic acid.

We assessed the effect of kynurenic acid on AS19-induced stimulatory activity of LC neurons in Wis (n = 6) and WKY (n = 8) rats by administrating cumulative doses of the glutamatergic antagonist (0.25-1 µmol i.c.v.) before the administration of AS19 (Fig. 2a-b). Repeated two-way ANOVA revealed that kynurenic acid potentiated the stimulatory effect induced by AS19 (1.25-10 mg/kg i.v.) on LC neurons from Wis rats (AS19 x Kyn: F_(3,44)_ = 4.22; p < 0.05; Fig. 2c), being statistically significantly for the last three doses (p < 0.01, Šídák’s multiple comparisons test). However, Kynurenic acid dampened the stimulatory effect of AS19 in WKY (AS19 x Kyn: F_(3,33)_ = 3.62, p < 0.05; repeated two-way ANOVA; Fig. 2d). Thus, this effect was much less pronounced compared to Wis rats (p < 0.05, unpaired two-sided Student’s t-test) (Fig. 2e), and it was not dependent on the kynurenic effect on the basal firing rate because the effect was not different in WKY compared to Wis (Fig. 2f).

**Figure 2.**
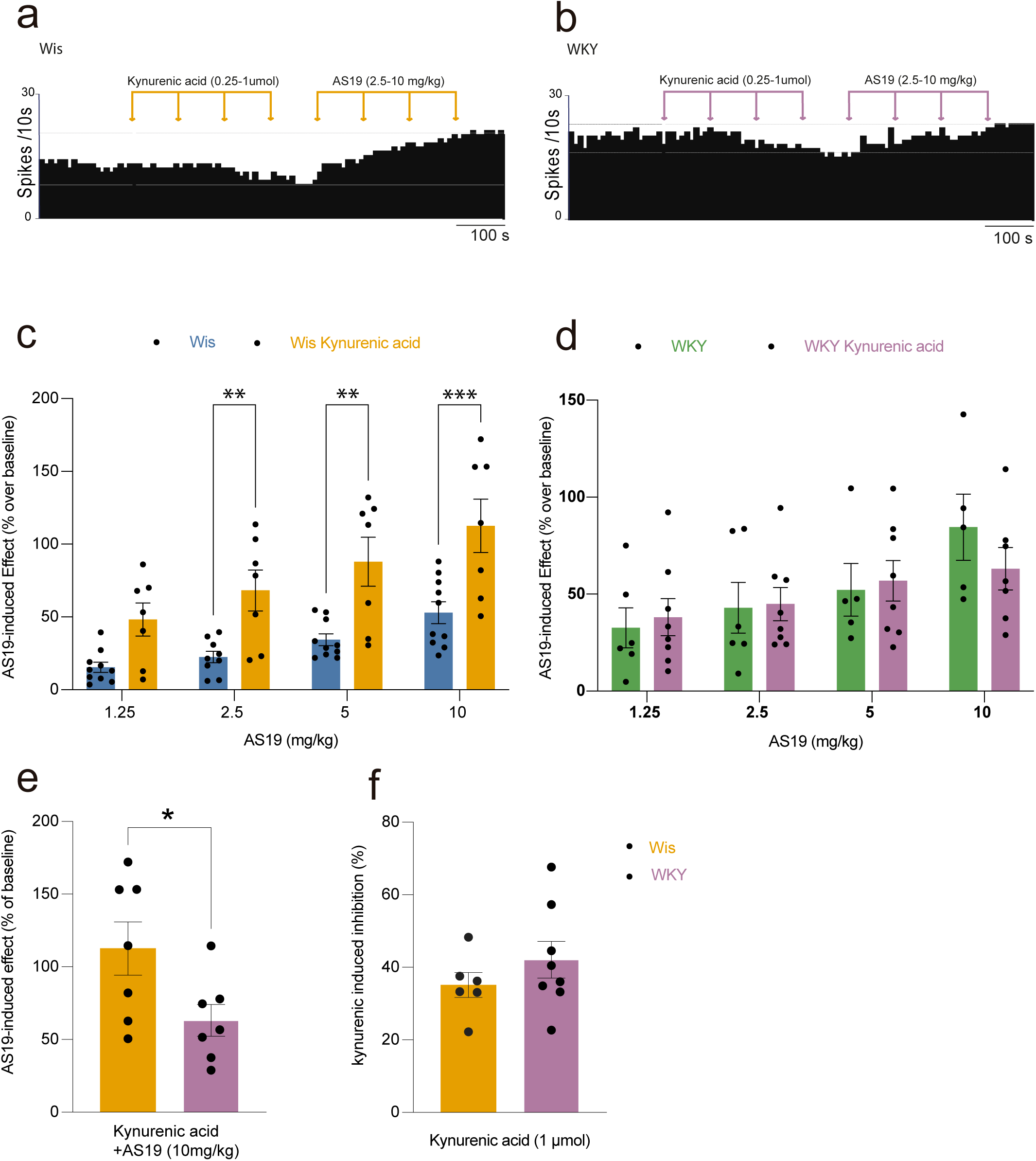
The stimulatory effect of AS19 on the basal firing rate of locus coeruleus (LC) noradrenergic neurons after kynurenic acid (KYN) administration in Wistar (Wis) and Wistar Kyoto (WKY) rats. (a-b) Representative firing rate histograms illustrate the stimulatory effect of AS19 (1.25–10 mg/kg, i.v.) on LC basal activity after kynurenic acid (0.25-1 µmol icv.) administration in Wis (a) and WKY rats (b). (c-d) Bar graphs showing kynurenic acid (1 µmol icv.)-induced effect on LC neurons basal activity and AS19 (10mg/kg, i.v.)-induced effect after kynurenic acid (1 µmol icv.) administration in Wis (n = 7 – 10; c) and WKY (n = 5 – 7; d) rats (**p < 0.01 and ***p < 0,001, Repeated two-way ANOVA). (e) Bar graphs comparing AS19 (10mg/kg)-induced effect after kynurenic acid administration in Wis and WKY rats. (f) Bar graphs showing kynurenic acid (1 µmol icv.)-induced inhibition on LC neurons basal activity in Wis (n = 7 - 10) and WKY (n = 5 - 7) rats. Bars represent mean ± S.E.M and circles individual cell values.

### 3.3. No differential in vitro effects of AS19 on the glutamatergic inputs in WKY rats

In order to further investigate the possible interaction of AS19 with the glutamatergic neurotransmission in the LC we recorded postsynaptic and presynaptic activity by performing in vitro patch-clamp assays. We identified LC neurons by the presence of a resting IRK conductance (see in Experimental Procedures).

Consistent with previous findings (Bruzos-Cidón et al., 2015), the spontaneous excitatory synaptic activity in the LC was not alter in the WKY rats, because neither the interevent interval nor the amplitude of sEPSC was different to Wis (Fig. 3a-c). Next, the effect of AS19 on spontaneous excitatory synaptic activity of LC neurons was recorded. For that purpose, we studied sEPSC interevent interval and amplitude before and during AS19 application (50 µM, 4 min) (Fig. 3a). AS19 perfusion significantly reduced presynaptic activity in both Wis (p < 0.01, paired two-tailed Student’s t-test; Fig. 3d) and WKY rats (p < 0.05, paired two-tailed Student’s t-test; Fig. 3e) in a similar extent (Fig. 3f). However, AS19 did not significantly alter the postsynaptic activity neither in Wis (Fig. 3g) nor in WKY rats (Fig. 3h) and therefore, no differences in the magnitude of the effect were found between strains (Fig. 3i). The evaluation of the kinetic parameters of sEPSC (rise time, decay time and half-width) revealed no differences between groups regarding the effect of AS19 (see Supplementary Materials: Table 1).

**Figure 3.**
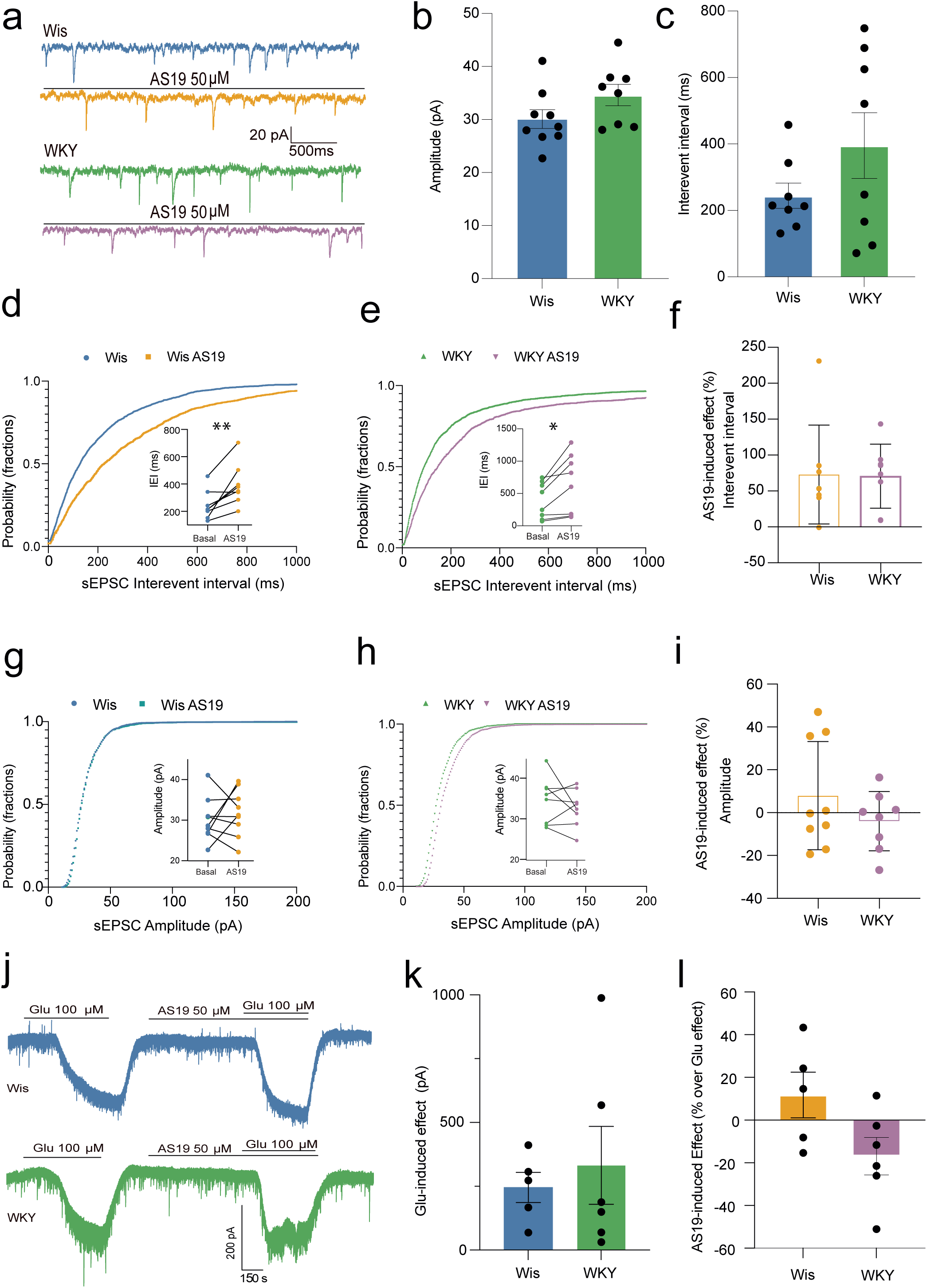
Glutamatergic transmission on locus coeruleus (LC) neurons in Wistar (Wis) and Wistar Kyoto (WKY) rats after AS19 administration. (a) Representative current traces of sEPSCs from LC neurons (n = number of neurons) before and during AS19 (50 µM) perfusion in Wis and WKY rats. (b) Bar graphs showing basal sEPSCs amplitude in Wis (n = 9) and WKY (n = 8) rats. (c) Bar graphs showing basal sEPSCs interevent interval in Wis (n = 8) and WKY (n = 8) rats. (d) Comparison of the inter-event interval distributions (IEI) of sEPSCs recorded from LC neurons of Wis (n = 8) rats under baseline conditions and following AS19 administration. Cumulative distribution functions of event amplitude for both conditions are shown. Inset shows mean IEIs before and after AS19 administration (** p < 0.01, paired student t-test). (e) Comparison of the inter-event interval distributions of sEPSCs recorded from LC neurons of WKY (n = 8) rats under baseline conditions and following AS19 administration. Cumulative distribution functions of event amplitude for both conditions are shown. Inset shows mean IEIs before and after AS19 administration (*p < 0.05, paired student t-test). (f) Comparison of the effect of AS19 on inter-event interval sEPSCs in LC neurons from Wis (n = 8) and WKY (n = 8) rats. The effect is expressed as a percentage change relative to baseline. (g) Comparison of amplitude distributions of sEPSCs recorded from LC neurons of Wis (n = 9) rats under baseline conditions and after AS19 administration. Cumulative distribution functions of event amplitude for both conditions are shown. Inset shows no significant difference in mean amplitude. (h) Comparison of amplitude distributions of sEPSCs recorded from LC neurons of WKY (n = 8) rats under baseline conditions and after AS19 administration. Cumulative distribution functions of event amplitude for both conditions are shown. Inset shows no significant difference in mean amplitude. (i) Comparison of the effect of AS19 on the amplitude of sEPSCs in LC neurons from Wis (n = 9) and WKY (n = 8) rats. The effect is expressed as a percentage change relative to baseline. Representative traces (j) of the inward current amplitude induced by a brief application of glutamate (100 μM) before and during AS19 (50 µM). The holding potential was -60 mV. (k) Bar graphs showing glutamate-induced inward current amplitude in Wis (n = 5) and WKY (n = 6) rats. (l) Bar graphs showing AS19-induced effect in percentage, over glu-induced inward current amplitude in Wis (n = 5) and WKY (n = 6) rats. Bars represent mean ± S.E.M and circles individual cell values.

**Table 1.** Kinetic parameters of spontaneous excitatory postsynaptic current (sEPSC) activity in LC neurons before and after AS19 administration.

|  | Wis<br>(n=8-9) | Wis AS19<br>(n=8-9) | WKY<br>(n=7-8) | WKY AS19<br>(n=7-8) |
| --- | --- | --- | --- | --- |
| sEPSC rise time<br>mean (ms) | 0.62 ± 0.04 | 0.74± 0.17 | 0.55± 0.04 | 0.62± 0.06 |
| sEPSC decay<br>time mean (ms) | 2.12± 0.08 | 2.11± 0.24 | 1.65± 0.19 | 1.97± 0.22 |
| sEPSC Half-width mean (ms) | 0.89± 0.07 | 0.89± 0.07 | 0.80± 0.07 | 0.89± 0.14 |

Wis: Wistar WKY: Wistar
Kyoto
All values represent the mean ± S.E.M. of n experiments.

Next, in a different set of experiments, the effect of AS19 on the inward current induced by L-glutamic acid (Glu) (100 µM) was evaluated in LC neurons (Fig. 3j). Thus, Glu caused an inward current of similar amplitude in both WKY and Wis rats (Fig.3k) and application of AS19 (50 µM) did not alter the amplitude of the Glu-induced current in either strain (Fig. 3l).

### 3.4. Altered pattern of 5-HT_7_ receptor expression in WKY rats

To further investigate if the in vivo observed differences induced by AS19 in LC neurons of WKY and Wis rats were due to differences in the 5-HT_7_ receptor expression, a western blot assay was performed in samples of several brain areas, including the LC, from Wis and WKY rats (Fig. 4a).

**Figure 4.**
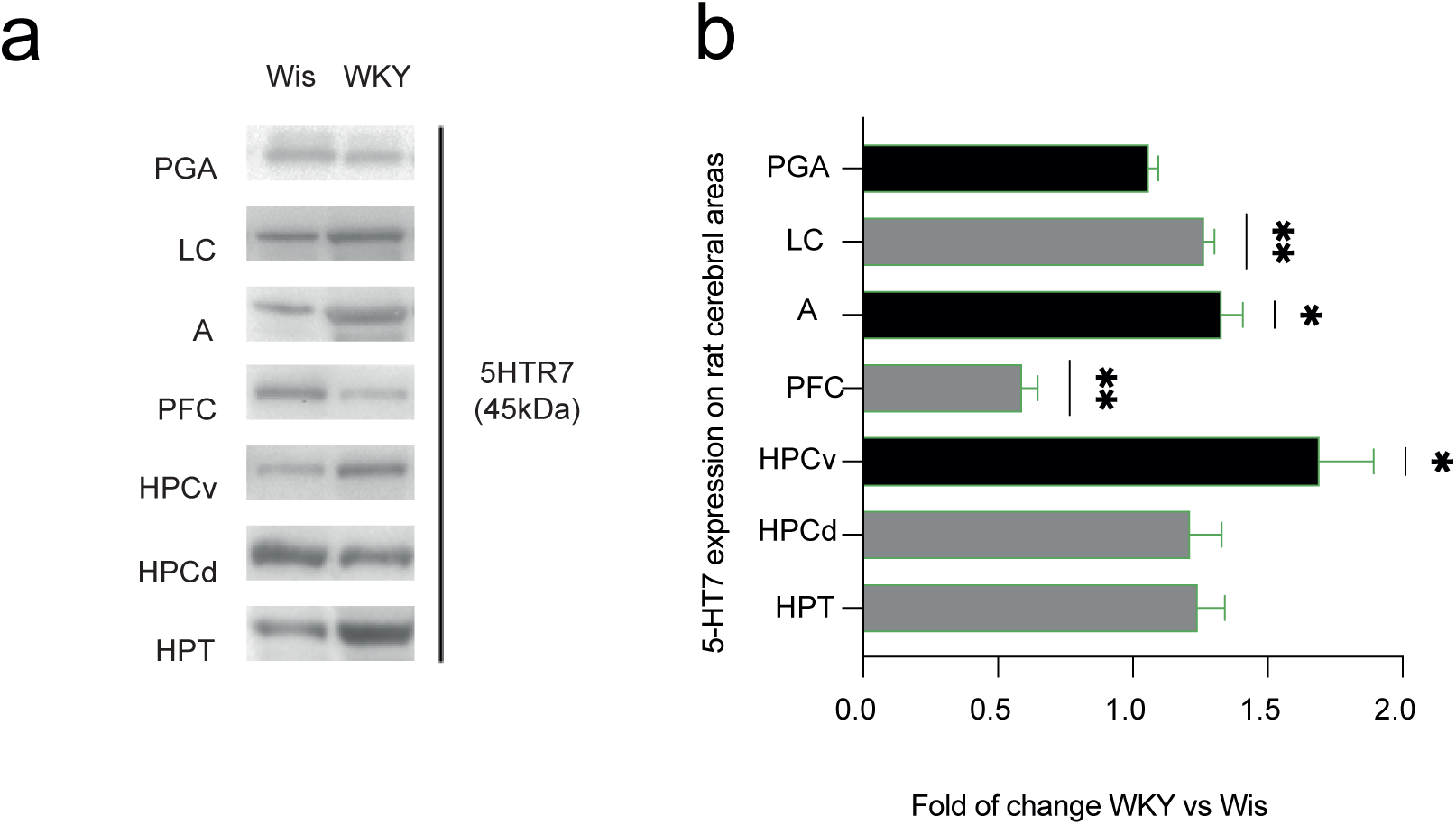
Brain 5-HT_7_ receptor expression in Wistar (Wis) and Wistar Kyoto (WKY) rats. (a) Western blot analysis of 5HTR7 expression in different brain regions of Wistar (Wis) and Wistar Kyoto (WKY) rats. (b) Bar graph showing the relative expression of 5-HT_7_ receptor protein in various brain regions, normalized to Wistar levels (Wis = 1.0), in both Wis (n = 4 - 5) and WKY (n = 4 - 5) rats. Bars represent mean ± S.E.M. Regions analyzed include the periaqueductal gray (PGA), locus coeruleus (LC), amygdala (A), prefrontal cortex (PFC), ventral hippocampus (HPCv), dorsal hippocampus (HPCd), and hypothalamus (HPT). Statistical analysis was performed using a one-sample Student’s *t*-test versus the normalized Wistar value (1.0) (*p < 0.05* *, p < 0.01 **).

HPT, HPCd, HPCv, PFC, A, PGA and LC nuclei were studied. Our data indicated that there were no statistical differences between strains in the 5-HT_7_ receptor expression on PGA, HPCd and HPT. However, 5-HT_7_ receptor expression was higher in WKY rats compared to that observed in Wis rats in LC, A and HPCv (p < 0.01 for LC and p < 0.05 for A and HPCv, One sample Student’s t-test). Finally, 5-HT_7_ expression appeared reduced in PFC from WKY rats compared to that in Wis rats (p < 0.01; One sample Student’s t-test) (Fig.4b).

## 4. Discussion

This study aimed to characterize the effects of serotonergic 5-HT_7_ receptor activation on LC function in WKY rats, a well-established model of treatment-resistant depression. In vivo and in vitro electrophysiological recordings were combined with receptor expression analyses in LC and related regions. In vivo, AS19, a 5-HT_7_ agonist, increased LC firing, with a greater effect in WKY than Wistar controls. Blocking glutamatergic transmission enhanced AS19 response in Wistar but not WKY rats. In contrast, AS19 did not significantly affect postsynaptic activity in LC slices, and presynaptic effects were similar across strains. Western blot revealed increased 5-HT_7_ expression in LC, HPCv, and A, but decreased expression in PFC of WKY rats.

WKY rats exhibit behavioral and neurobiological traits consistent with treatment-resistant depression, including Hypothalamic–Pituitary–Adrenal axis dysregulation, altered monoaminergic and glutamatergic signaling, neuroinflammation, and impaired neurotrophic support (Aleksandrova et al., 2019; Li et al., 2022; Millard et al., 2020; Redei et al., 2023). Consistent with previous findings, this strain shows heightened noradrenergic activity, reflected in increased LC firing and burst rates (Bruzos-Cidón et al., 2014; El Mansari et al., 2023). AS19 enhanced LC firing in both strains, with a markedly greater effect in WKY rats. This response was prevented by SB-269970, confirming receptor-specific mediation.

The activation of 5-HT_7_ receptors influences noradrenergic neurotransmission indirectly via serotonergic projections from the DRN (Kusek et al., 2015). These receptors are highly expressed in DRN GABAergic interneurons, where activation increases inhibitory tone and reduces serotonin release (Monti & Jantos, 2006). Because the LC receives tonic serotonergic inhibition (Haddjeri et al., 1997), 5-HT_7_ activation in the DRN may decrease serotonergic output and disinhibit LC neurons. In fact, AS19 reduces DRN activity in vivo, (Mnie-Filali et al., 2011). Thus, the absence of AS19 effects in LC slices supports this interpretation, as in vitro preparations lack raphe inputs, suggesting that the observed modulation depends on circuit-level connectivity rather than direct LC receptor activation. Although previous work assessed AS19 in vivo at systemic doses, the concentration used here (50 µM) ensures effective activation in slices, where diffusion barriers require higher local doses. This approach is consistent with electrophysiological protocols for serotonergic agonists, which typically employ concentrations in the 10–50 µM range to achieve maximal GPCR activation in slices (Haddjeri et al., 1998; Lucas et al., 2007).

The greater AS19 response in WKY rats aligns with reduced inhibitory regulation within the LC. Previous work shows desensitization of 5-HT_1A_ and α_2_ receptors (Bruzos-Cidón et al., 2014) and altered GABAergic signaling (Bruzos-Cidón et al., 2015). These changes, plus impaired GABAergic function in the amygdala (Jiao et al., 2011), habenula (Korlatowicz et al., 2023), and cortical circuits (Inavally & Sadananda, 2024), may contribute to chronic noradrenergic hyperactivity in WKY rats.

Our analyses revealed elevated 5-HT_7_ levels in LC of WKY rats. Postsynaptic presence remains uncertain (Vanhoenacker et al., 2000); detected signal may reflect presynaptic serotonergic terminals from DRN (Kusek et al., 2015; Roberts et al., 2004). Increased receptor expression could enhance serotonergic control of LC inhibition. Conversely, reduced 5-HT_7_ in PFC may weaken top-down LC modulation. Although direct evidence for LC–PFC linkage is limited, 5-HT_7_ activation regulates glutamatergic output from PFC to DRN (Harsing, 2006; Olusakin et al., 2020; Pehrson & Sanchez, 2014). Thus, 5-HT_7_ receptor-mediated inhibition of glutamate release in DRN could reduce 5-HT output to LC, influencing noradrenergic tone.

When glutamatergic transmission was blocked, AS19-induced LC firing increase was smaller in WKY than Wis rats, suggesting WKY LC, already hyperactive and inhibition-deficient, has limited capacity for additional excitation. In contrast, Wis LC shows greater modulatory range. This blunted responsiveness aligns with reduced sensitivity to monoaminergic antidepressants (Bruzos-Cidón et al., 2015; Lahmame et al., 1997; Li et al., 2022), supporting circuit-level rigidity as a mechanism for treatment resistance.

Enhanced 5-HT_7_ expression in vHPC and A of WKY rats may further contribute to depressive phenotypes. Chronic stress upregulates 5-HT_7_ in hippocampal CA1, activating MMP-9 pathways that alter dendritic spines and reduce plasticity (Bijata et al., 2022; Naumenko et al., 2014). WKY rats show deficits in long-term potentiation at Schaffer collateral–CA1 synapses (Aleksandrova et al., 2020) and decreased synaptic protein expression, correlating with depressive-like behaviors (Li et al., 2022). In A, 5-HT_7_ modulates inhibitory control over principal neurons (Kusek et al., 2021), and dysregulation may underlie emotional disturbances in TRD (Fukui et al., 2025). Overexpression also affects PFC-to-DRN circuit maturation, leading to long-term depressive-like behaviors (Olusakin et al., 2020). Together, these findings support that enhanced 5-HT_7_ signaling in limbic structures perpetuates depression, whereas receptor blockade or deletion exerts antidepressant-like effects (Bijata et al., 2022; Gottlieb et al., 2023).

This study is limited by the use of a single agonist (AS19), potential off-target effects on 5-HT_1A_ or α_2_ receptors, and the absence of behavioral data linking LC modulation to antidepressant outcomes. However, AS19 was chosen because systemic administration at comparable doses suppresses DRN serotonergic activity (Mnie-Filali et al., 2011), supporting its relevance as a 5-HT_7_ probe. Future studies should integrate electrophysiology with behavioral paradigms and region-specific manipulations to clarify causal links between 5-HT7 signaling, LC dynamics, and treatment response.

In summary, 5-HT_7_ receptor activation exerts complex, state-dependent modulation of LC activity that depends on the integrity of glutamatergic and serotonergic circuits. The hyperactive and inhibitory-deficient state of the WKY LC limits its responsiveness to serotonergic modulation, potentially explaining resistance to monoaminergic antidepressants. Elevated 5-HT_7_ receptor expression in the HPC and A further supports its involvement in depressive-like behaviors and positions this animal model of DRT, and in particular the 5-HT_7_ receptor signaling, as a promising target to investigate the etiology and develop novel therapeutic approaches for depression resistant to monoaminergic pharmacological treatments.

## 5. Conflict of Interest

The authors declare that the research was conducted in the absence of any commercial or financial relationships that could be construed as a potential conflict of interest.

## 6. Author Contributions

Conceptualization, L.U and M.T; methodology, C.B.-C. and N.L; validation, M.T., JV-L. and L.U; formal analysis, C.B.-C. and N.L; investigation, C.B.-C and H.B; resources, M.T. and C.B.-C.; data curation, C.B.-C. and N.L.; writing— original draft preparation, C.B.-C. and M.T; writing—review and editing, C.B-C, M.T, N.L, J.V-L, H.B; visualization, C.B-C, N.L, H.B; supervision, M.T. and J.V-L; revision and correction of the article, C.B.-C, N.L. M.T. All authors have read and agreed to the published version of the manuscript.

## 7. Funding

This work was supported by grants from the University of the Basque Country, UPV/EHU (UFI 11/32); and the Spanish Government (FIS PI12/00613). CB-C and N-L had a predoctoral fellowship from the University of the Basque Country (UPV/EHU). These institutions had no further role in the present study. The authors were entirely responsible for the scientific content of the manuscript and the decision to submit it for publication. We thank The Advanced Research Facilities (SGIker) of the University of the Basque Country, UPV/EHU for animal care.

## Data Availability Statement

Relevant experimental details necessary for reproducibility have been indicated in the Materials and Methods section and can be provided to interested researchers upon request. All protocols, analytic methods, and study materials used in this research are available upon reasonable request from the corresponding author.

## Notes

### Competing Interest Statement

The authors have declared no competing interest.

